# A versatile high throughput screening platform for plant metabolic engineering highlights the major role of *ABI3* in lipid metabolism regulation

**DOI:** 10.1101/853606

**Authors:** Benjamin Pouvreau, Cheryl Blundell, Harpreet Vohra, Alexander B. Zwart, Taj Arndell, Surinder Singh, Thomas Vanhercke

## Abstract

Traditional functional genetic studies in crops are time-consuming, complicated and cannot be readily scaled up. The reason is that mutant or transformed crops need to be generated to study the effect of gene modifications on specific traits of interest. However, many crop species have a complex genome and a long generation time. As a result, it usually takes several months to over a year to obtain desired mutants or transgenic plants, which represents a significant bottleneck in the development of new crop varieties.

To overcome this major issue, we are currently establishing a versatile plant genetic screening platform, amenable to high throughput screening in almost any crop species, with a unique workflow. This platform combines protoplast transformation and fluorescence-activated cell sorting.

Here we show that tobacco protoplasts can accumulate high levels of lipids if transiently transformed with genes involved in lipid biosynthesis and can be sorted based on lipid content. Hence, protoplasts can be used as a predictive tool for plant lipid engineering. Using this newly established strategy, we demonstrate the major role of *ABI3* in plant lipid accumulation.

We anticipate that this workflow can be applied to numerous highly valuable metabolic traits other than storage lipid accumulation. This new strategy represents a significant step towards screening complex genetic libraries, in a single experiment and in a matter of days, as opposed to years by conventional means.

## Introduction

Modern societies have slowly transitioned, over the past couple of centuries, from a bio-based economy to a petrol-based economy. Agriculture consequently transitioned to a food/feed focus whereas it used to meet all demands for energy and materials too. Driven by the threat of climate change and a need for more sustainability, re-establishing a bio-based economy is currently a major global challenge. To compete with established petrochemical industries, crops need to be re-engineered to have new traits or produce new compounds. However, current crop metabolic engineering approaches are slow and difficult, and this creates a significant bottleneck in the development of new crop varieties that are tailored to meet future agricultural needs in an ever-changing environment.

As reflected by the late Richard Feynman’s statement: “What I cannot create, I do not understand”, the major challenge in crop metabolic engineering is to find the right set of genetic components necessary to obtain a desired trait. This is particularly due to traditional functional genetic studies in crops being time-consuming, complicated and not usually compatible with high throughput. Indeed, it usually takes several months to over a year to obtain desired mutants or transgenic plants (Ishida et al., 2007; Yadava et al., 2017) necessary to study the effect of single genetic components on specific traits of interest. The testing of complex metabolic or genetic circuits thus seems like a herculean task. The use of model plants, such as *Arabidopsis thaliana*, with fully annotated genomes and much faster and cheaper gene delivery systems (Clough and Bent, 1998), had an undeniable impact on crop improvement (Provart et al., 2016), but still has limitations regarding time, scale and technology transfer between species.

While more promising, few transient transformation systems have been developed to test genetic components for crops re-engineering strategies. Currently, the most widely used transient transformation system for rapid gene testing is agroinfiltration of *Nicotiana benthamiana* leaves (Yang et al., 2000), which allows for the delivery of one or multiple genetic components to test multigene circuits in a few days (Wood et al., 2009). Pioneering studies have already demonstrated the power of such efficient transient transformation systems. For example, the ‘Leaf Oil’ platform technology, which allows plants to accumulate oil in all vegetative organs (Vanhercke et al., 2017) with yields potentially exceeding those of canola and oil palm, was quickly developed thanks to an extensive number of design-built-test-learn cycles (El Tahchy et al., 2017; Vanhercke et al., 2014; Vanhercke et al., 2013) that strongly relied on an efficient use of the agroinfiltration-based system. The technology is continuously being upgraded by further cycles to create tailor-made products for biofuel applications (Reynolds et al., 2017; Reynolds et al., 2015), and is also being transferred to a range of major crops such as sugarcane (Zale et al., 2016), maize (Alameldin et al., 2017) and *Sorghum bicolor* (Vanhercke et al., 2019). Such high biomass crops have the potential to become a future sustainable and abundant supply of plant oils for food, feed, biofuel and oleo-chemical applications (Rahman et al., 2016; Weselake, 2016). This type of assay is not limited to studying leaf specific metabolic pathways. Petrie et al. demonstrated that co-infiltration of the tested components in combination with the *Arabidopsis thaliana LEAFY COTYLEDON2* (*LEC2*) gene results in a leaf with a seed-like gene regulation profile, thus allowing for the testing of seed-specific components (Petrie et al., 2010). A modified version of this assay was also developed in transgenic *LEC2* induced *Brassica napus* somatic embryos for testing seed-specific components with increased reliability to predict seed traits (Belide et al., 2013). These assays allowed for the rapid gene testing and assembly of the ω3 fish-like oil synthetic seed biosynthesis pathway that was first deployed in *Arabidopsis thaliana* as proof-of-concept (Petrie et al., 2012), and then in oilseed crops such as *Camelina sativa* (Petrie et al., 2014) and canola (Walsh et al., 2016), with ω3 production at industrially-relevant levels. Nevertheless, these transient transformation systems are still limited in terms of scale, species and tissue specificity, and other transient assays are required to overcome these limitations.

Another type of transient system based on protoplast transfection has been available for more than 50 years, and now represents a very promising system for rapid testing of genetic components in most plants, and for several reasons. First, protoplasts can be isolated from almost any tissue of any crop (Burris et al., 2016), thus allowing better predictability through more species-targeted approaches than the agroinfiltration-based system that is established only in tobacco and few other species. Second, protoplasts are single cells, thus allowing more precise studies than multicellular systems to address tissue and cell-type specific questions. Thirdly, a very large number of protoplasts can be isolated from one preparation, allowing testing of many variables simultaneously in one assay. Finally, protoplast transformation is now well established (Yoo et al., 2007), with a higher efficiency than any other plant transformation system (Jiang et al., 2013), and is therefore very-well suited for high throughput platforms. The use of protoplast-based systems also allows access to other types of high throughput approaches, such as flow cytometry and cell sorting, which open a wide new range of screening possibilities. Protoplasts have long been extensively used with fluorescence-activated cell sorting for several different applications. Such methods were developed from the early 80’s for applications such as protoplast viral infection level monitoring (van Klaveren et al., 1983) and chlorophyll content and size distribution characterization (Galbraith et al., 1988). More recently, similar approaches have been used for more complex studies such as protein accumulation detection (Cronjé et al., 2004), overexpression analyses (Bargmann and Birnbaum, 2009), sorting of particular cell types for RNA isolation and transcriptome-wide analyses (Bargmann and Birnbaum, 2010; Birnbaum et al., 2005), or to study cell-type specific expression during both development and stress (Grønlund et al., 2012). Such techniques allow for the direct screening of millions of variants in a very short amount of time, and ongoing improvements (You et al., 2014) allow for the development of increasingly powerful high throughput screening methods. For example, recently, cell sorting based on the lipid content of thousands of mutant lines of a green unicellular algae was achieved in a few hours, followed by a few additional days for cellular multiplication and additional lipid analyses to confirm the phenotype (Terashima et al., 2015), representing one of the most powerful mutant screens ever reported for green vegetal cells. While random mutagenesis is not available for protoplasts, transient transformation is very efficient, and transformation of expression libraries could be carried out to generate large numbers of transformed cells for rapid trait evaluation by flow cytometry. However, sorting of plant protoplasts based on a high-value trait, such as lipid accumulation, has not yet been reported.

The regulation of lipid biosynthesis and storage in plants has been extensively studied. The highly conserved plant transcription factors ABSCISIC ACID INSENSITIVE 3 (ABI3) (Giraudat et al., 1992), FUSCA3 (FUS3) (Luerssen et al., 1998), LEAFY COTYLEDON1 (LEC1) (Meinke et al., 1994; West et al., 1994), and LEC2 (Chen et al., 1997) are the major master regulators that control the gene regulation networks governing most seed developmental mechanisms (Braybrook et al., 2006; North et al., 2010; Suzuki et al., 2003; To et al., 2006; Wang and Perry, 2013). The *abi3, fus3, lec1* and *lec2* mutants share common phenotypes such as reduced storage protein accumulation and anthocyanin accumulation (To et al., 2006). FUS3, LEC1 and LEC2 also have been reported to trigger triacylglycerol accumulation in seeds, leaves and liquid cell culture (Lotan et al., 1998; Santos Mendoza et al., 2005; Tan et al., 2011; Vanhercke et al., 2017; Zhang et al., 2016; Zhang et al., 2002). The major action of LEC1, LEC2 and FUS3 on lipid accumulation is mostly through regulating directly or indirectly the expression of the Wrinkled1 (WRI1) transcription factor (Baud et al., 2007; Casson and Lindsey, 2006; Marchive et al., 2014; Santos-Mendoza et al., 2008; Wang and Perry, 2013; Yamamoto et al., 2010), but the complete picture of *WRI1* regulation is still unclear (Kim et al., 2016). *WRI1* function is highly conserved amongst plants. *WRI1* is a node acting at the interplay between these master regulatory elements and fatty acid metabolism. *WRI1* directly regulates the transcription level of many genes essential for fatty acid synthesis to trigger seed oil production (Baud et al., 2007; Pouvreau et al., 2011). After it was demonstrated that *WRI1* overexpression increases oil accumulation without undesirable side effects (Pouvreau et al., 2011; Shen et al., 2010), *WRI1* became a major target of metabolic engineering approaches to increase oil content (Vanhercke et al., 2017; Vanhercke et al., 2014; Vanhercke et al., 2013). By contrast, despite the fact that many of ABI3 targeted genes are known to function in seed oil storage (Monke et al., 2012; Wang et al., 2007), to date, little is known about the direct effect of *ABI3* on lipid content in plants.

In this study, we demonstrate that protoplasts from 15-day-old tobacco leaves protoplasts can accumulate high levels of lipids if transiently transformed with genes involved in lipid biosynthesis. We observed a direct correlation between protoplast lipid content and protoplast fluorescence intensity after lipid-specific fluorescent staining. Furthermore, we show that protoplasts can be sorted based on lipid content and further used for downstream processing and analyses. Transient gene expression in protoplasts produced metabolic results similar to those reported for stably transformed plants, in terms of lipid accumulation. We demonstrate that combinatorial gene testing is possible in protoplasts and we observed a similar synergistic effect of the genes *WRI1* and *DGAT1* on increasing lipid accumulation in protoplasts, compared with previously published studies conducted in transiently and stably transformed leaves (Vanhercke et al., 2014; Vanhercke et al., 2013). Taken together, our results demonstrate that tobacco leaf protoplasts are a reliable model for plant lipid engineering. This platform constitutes a new high throughput tool to develop plant lipid engineering strategies, but also an alternative strategy to study plant lipid metabolism in general, such as the relative effects of major known elicitors of lipid accumulation in plants. We compared the effect of *ABI3, FUS3, LEC1* and *LEC2* on lipid accumulation in protoplasts. Our results confirm that *ABI3* is an important regulator of oil accumulation and demonstrate for the first time the direct correlation between *ABI3* overexpression and increased lipid content in plants. These results also suggest that *ABI3* might trigger lipid accumulation partly through pathways independent from *WRI1* and *LEC2* regulation.

## Material & Methods

### Plant material and growth conditions

Plants used in this study all originated from the *Nicotiana tabacum* cultivar Wisconsin 38, including wild-type (WT) and a T4 generation *WRI1*-*DGAT1*-*OLEOSIN* transgenic line, referred to as HO (High-Oil), for the high levels of triglycerides accumulating in its leaves (Vanhercke et al., 2014). Seeds were surface-sterilised by 2 min incubation in 70% ethanol, followed by 10 min incubation in 1.8% bleach and five washes in sterile water, all with frequent agitation. Seeds were sown on half MS media, supplemented with 3% sucrose, in petri dishes and placed in the dark at 4 °C for 4 days. Plants were grown in a growth cabinet under 16 h day/8 h night light conditions (100 μmol m^−2^ s^−1^) at 22 °C/18 °C, respectively.

### Plasmids DNA construction and preparation

Vectors were designed as minimal plant expression constructs for production in large quantity in *E. coli* and strong transient expression in tobacco protoplasts. A modified pENTR11 vector (Invitrogen) was used as a minimal backbone after removal of the ccdB operon, conserving only kanamycin resistance and the *E. coli* origin of replication. The chemically synthesised (GeneArt) *CaMV35S* promoter, coding sequence (Arabidopsis thaliana WRI1, DGAT1, ABI3, FUS3, LEC1 or LEC2, respectively AT3G54320.1, AT2G19450.1, AT3G24650.1, AT3G26790.1, AT1G21970.1 or AT1G28300, codon optimised for tobacco) and NOS terminator were ligated in this minimal vector. For protoplast transient transformation, plasmid DNA was prepared using the NucleoBond Xtra Midi Kit (MACHEREY-NAGEL, Germany) according to the manufacturer’s recommended protocol, and resuspended in ultrapure DNAse/RNAse-free distilled water (Invitrogen) at 1 μg/μl.

### Protoplasts preparation, transient transformation and staining

The whole procedure was conducted under sterile conditions. Plants were collected 15 days after germination. Leaf tissues were harvested and thinly sliced in protoplast buffer (400mM mannitol, 10mM MES hydrate, 20mM KCl, 20mM CaCl2, 2mM MgCl2, pH 5.7, 0.1% bovine serum albumin). Leaf strips were rinsed with fresh protoplast buffer, placed in enzyme solution (protoplast buffer supplemented with 1.2% cellulase Onozuka R-10, 0.6% cellulase Onozuka RS, and 0.4% macerozyme R-10), subjected to vacuum for 10 min and incubated in enzyme solution for 16h in the dark at 22 °C. After incubation, protoplasts were released with 5 min agitation on a rotating plate at 50rpm, filtered through a 40μm cell strainer, washed twice with fresh protoplast buffer, counted on a hemocytometer, and then resuspended in fresh protoplast buffer at 0.25M cells per ml. For transformation, protoplast samples were incubated for 1h on ice, the buffer was then removed, pelleted protoplasts were resuspended in an identical volume of MMg solution (0.4M mannitol, 15mM MgCl2, 4mM MES, pH 5.6), supplemented with 25μg/ml of each plasmid and then mixed with an identical volume of PEG solution (40% PEG4000, 0.2M mannitol, 300mM CaCl2). After 5 min incubation, protoplasts were washed twice with fresh protoplast buffer, resuspended in fresh protoplast buffer at 0.25M cells per ml, placed in culture plates and incubated in a growth cabinet under constant light conditions (150 μmol m^−2^ s^−1^) at 22 °C. For lipid staining, protoplasts in protoplast buffer at 0.25M cells per ml were supplemented with 1 μg/ml BODIPY™ 493/503, incubated for 15 min at room temperature in the dark, spun down (300 x g for 5 min) and resuspended in the same volume of fresh protoplast buffer.

### Lipid extractions and analyses

For total lipid extractions, fresh leaves were cut as for protoplast isolation and freeze-dried whereas freshly isolated protoplasts were spun down (for 5 min) to remove all buffer and then snap-frozen in liquid nitrogen. Samples were ground with a ball bearing in TissuLyser II (QIAGEN, Germany) and homogenised in 36% methanol, 36% chloroform, 20 mM acetic acid, 1mM EDTA. Total lipid fraction (lower phase) was separated by 2 min centrifugation at 4000 x g, dried under nitrogen and resuspended in chloroform. Following thin layer chromatography fractionation of total lipid extracts, fatty acid methyl ester quantification and analysis by gas chromatography were conducted as previously described (El Tahchy et al., 2017; Vanhercke et al., 2013). Unless otherwise stated, analyses were conducted on a minimum of three individual samples for each genotype, condition or cell population.

### Cell imaging by microscopy

Glass slides were prepared by placing four drops of petroleum jelly in a square arrangement and 50ul of protoplasts at 1M cells per ml in protoplast buffer within that square. A coverslip was then gently pressed down on the solution and jelly drops. Protoplasts were captured (**figure 3A**) using a 40× objective on a Zeiss Axio Imager M1 fluorescence compound microscope fitted with an AxioCam HR Rev3 camera (Carl Zeiss Microscopy GmbH, Germany), with bright field and no filter or with U.V. light and a Filter Set 38 HE (Excitation filter BP 450-490nm, Beam splitter FT 495 HE, emission BP 500-550nm) to visualize lipid bodies stained with BODIPY™ 493/503. Images were acquired using the Zeiss Zen imaging software (blue edition, Carl Zeiss Microscopy GmbH, Germany).

**Figure 1.**
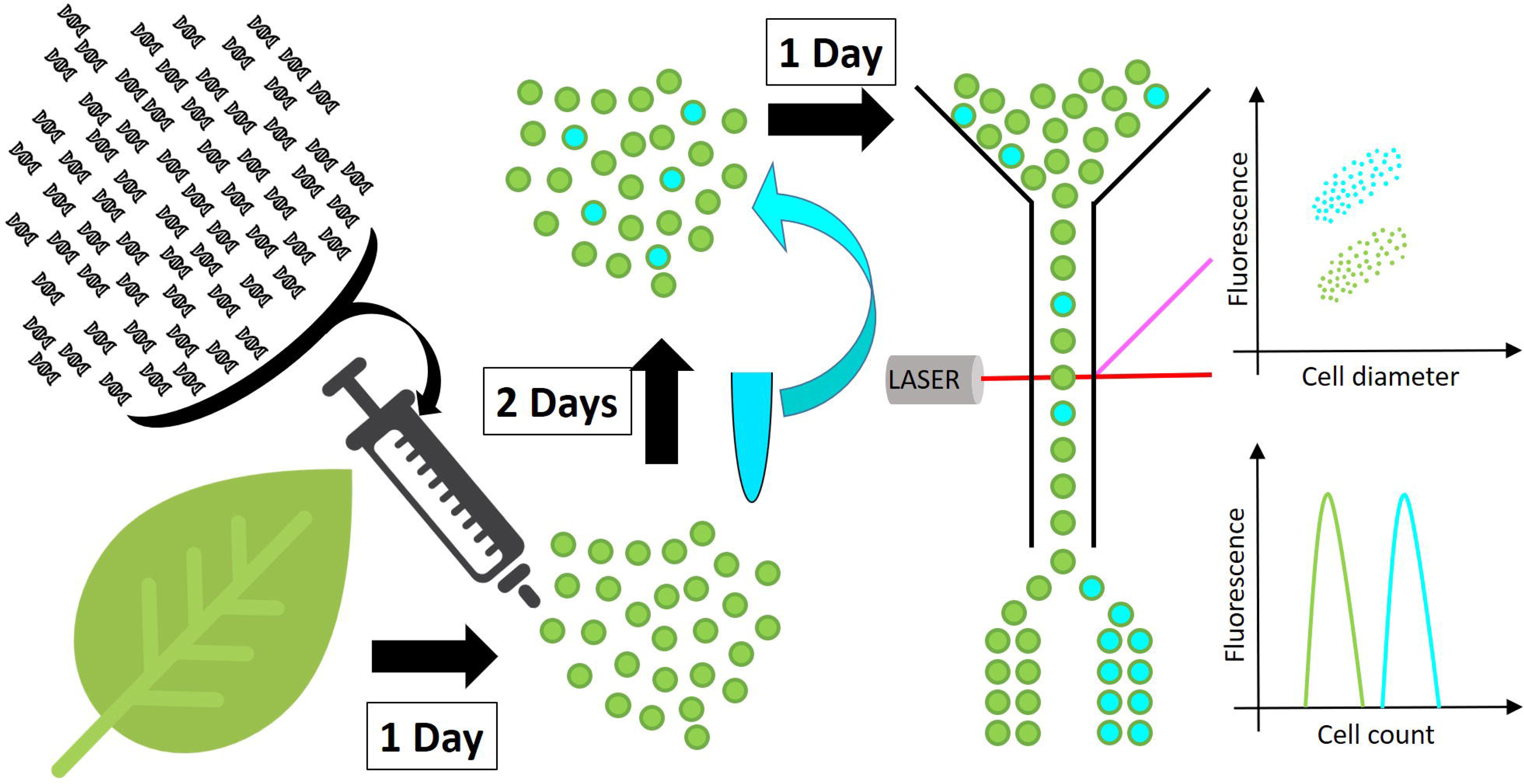
Schematic representation of the high throughput screen developed in this work. Protoplasts (green spheres) are isolated from a leaf and transformed with DNA constructs. After 48h incubation, to allow for transgene expression and metabolite accumulation, protoplasts are stained with a neutral lipid-specific fluorescent stain and screened for fluorescence by flow cytometry or fluorescence-activated cell sorting.

**Figure 2.**
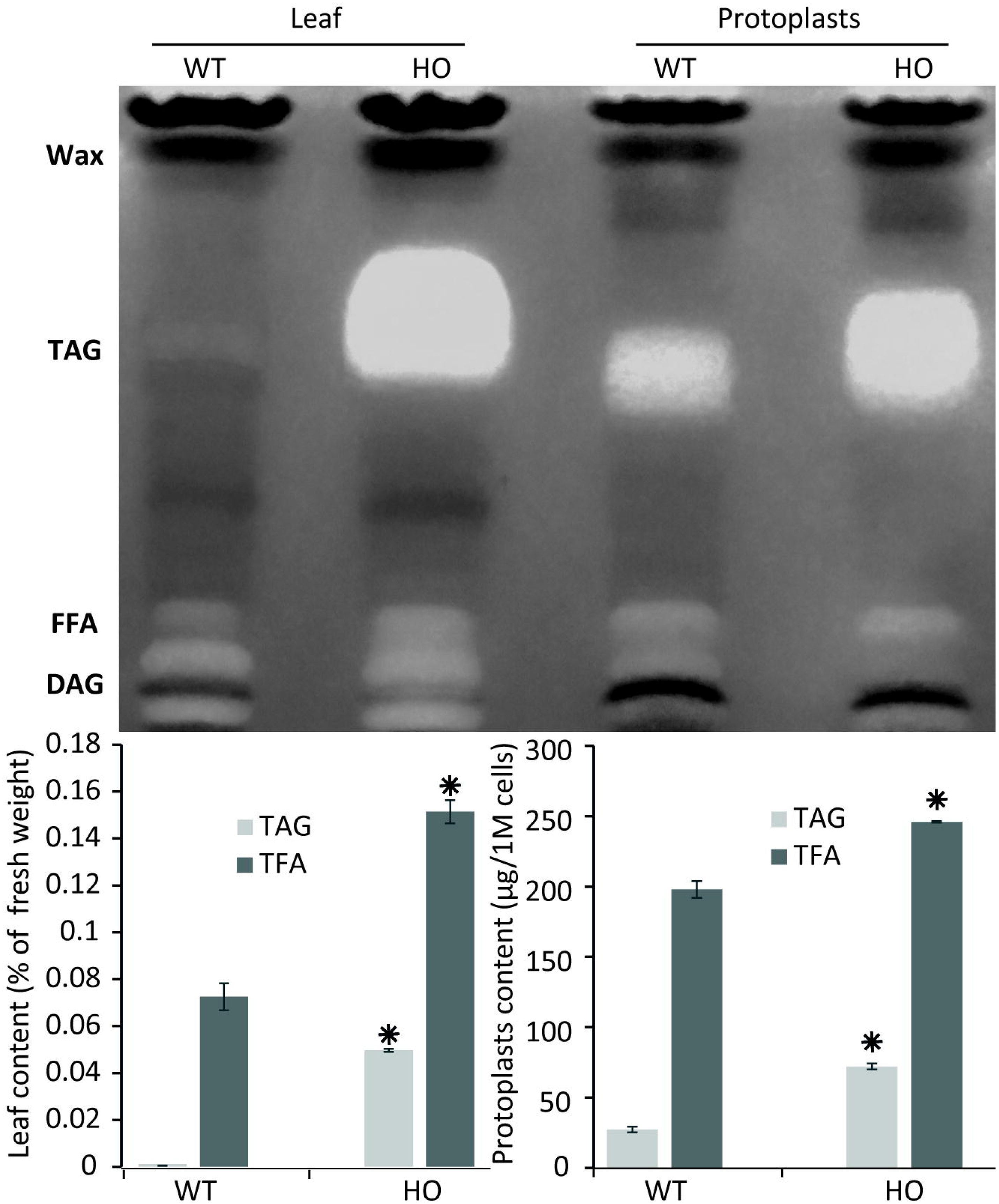
Protoplast lipid quantification results mimic total tissue lipid analyses results. Thin-layer-chromatography plate separation of total fatty acids (TFA) extracted from leaves or from isolated protoplasts of WT or HO lines and total lipid and triacylglycerol (TAG) quantification of the same samples quantified by gas chromatography. Concentrations are given in percent relative to leaf fresh weight or in micrograms per million cells. Bars represent standard deviation. Stars indicate statistically relevant variation compared to WT (P values < 0.05). FFA = free fatty acids, DAG = diacylglycerol.

**Figure 3.**
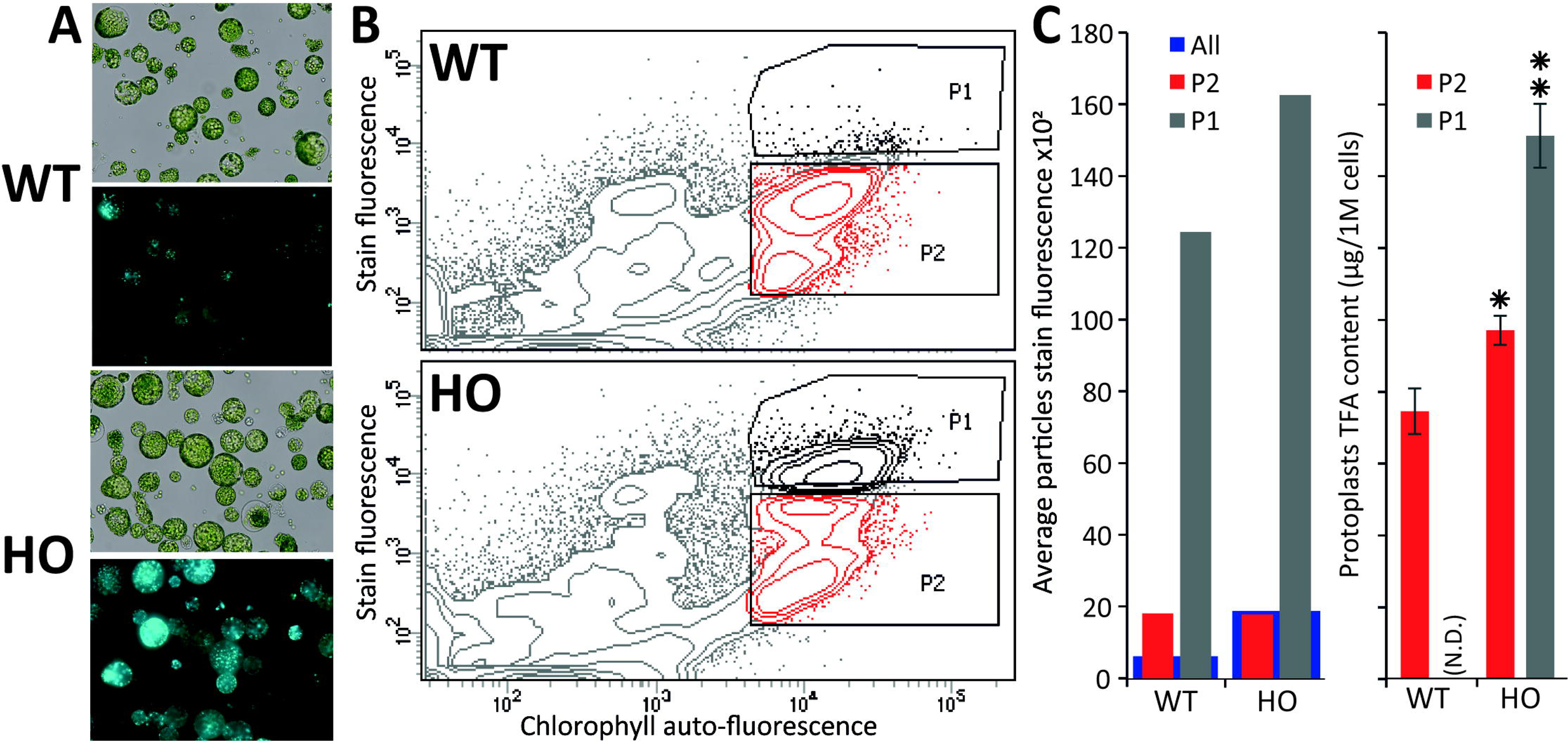
Fluorescence-activated cell sorting of lipid rich tobacco leaf protoplasts. **(A)**, Microscopy images of protoplasts extracted from WT or HO (line with high leaf oil content) tobacco leaves in bright field (top) and their lipid content (bottom), visualized under fluorescent light after BODIPY™ 493/503 staining. **(B)**, Cell sorting of the same protoplast samples, with the P1 population defined as the protoplasts with the highest stain fluorescence to chlorophyll auto-fluorescence ratio, and P2 defined as the remainder of the intact cells (based on previous data not show). **(C)**, Average stain fluorescence (left) and lipid content (right) for each sorted population. Average stain fluorescence of each whole sample is also indicated (blue bars). Where N.D. is in brackets, this indicates not determined. Error bars represent the standard deviation. Asterisks represent statistically significant differences (P value<0.05) compared with WT (one asterisk) or with both WT and HO P2 population (two asterisks).

### Protoplast flow cytometry and fluorescence-activated cell sorting

Cell sorting was performed on a BD FACS Aria III (BDIS, San Jose) with a 100 μm nozzle and data were acquired and analysed using the DIVA 8.0 software. Protoplasts suspended in the protoplast buffer at a concentration of 1 million/ml were acquired at an approximate flow rate of 10ul/min with mild agitation. Data were acquired for 30,000 particles and after exclusion of doublets, the samples were analysed for BODIPY™ 493/503 (Ex. 488nm, Em. 530/30 nm) and chlorophyll auto fluorescence (Ex. 640nm, Em. 670/30nm). Cells were sorted based on BODIPY/chlorophyll signal ratio. For each gated population, three independent samples of 0.25M cells were collected in protoplast buffer.

Flow cytometric analyses were conducted on an Invitrogen Attune NxT Flow Cytometer and analysed with Invitrogen Attune NxT Software. Cells suspended in the protoplast buffer at a concentration of 0.25M cells per ml were analysed at a flow rate of 250 μl/min and data were acquired for 300 μl. BODIPY™ 493/503 fluorescence was excited with a 488-nm laser and emission was captured by a 530/ 30-nm bandpass filter. Unless otherwise stated, analyses were conducted on a minimum of three individual samples for each condition.

### Statistical analysis

For the statistical analysis presented in figure 5, data were analysed via a linear mixed model fitted using the ‘lmer’ linear mixed modelling function from the ‘lme4’ package (Bates et al., 2015) of the R statistical software system (Team, 2014). This allowed the analysis to properly account for the impacts of the grouping of processed samples together on plates (via a factor ‘Plate’), and also recognise that the full dataset is formed from two individual repeats of the same experiment (via a factor ‘Experiment’). Hence, the model incorporated a (random effects) ‘blocking’ structure consisting of ‘Sample, nested within Plate, nested within ‘Experiment’. Treatment (fixed effects) structure was modelled as a 2-way factorial, *WRI1* crossed with *DGAT1*, where factors *WRI1* (having levels WRI1+ and WRI1-) and *DGAT1* (levels DGAT1+ and DGAT1-) represent the presence/absence of the corresponding genes in the samples. Hence, the WT treatment corresponds to the combination ‘WRI1-:DGAT1-’ and the samples transformed with *DGAT*1, *WRI1* or both, correspond to ‘WRI1-:DGAT1+’, ‘WRI1+:DGAT1-’ and ‘WRI1+:DGAT1+’ respectively. The model allows to test for the ‘main effect’ of *WRI1* (the average effect of *WRI1* status, regardless of *DGAT1* status), the main effect of *DGAT1* (the average effect of *DGAT1* status, regardless of *WRI1* status), and the interaction effect between *WRI1* and *DGAT1* (the combined effects of *WRI1* and *DGAT1* after having removed effects of the two main effects terms).

**Figure 4.**
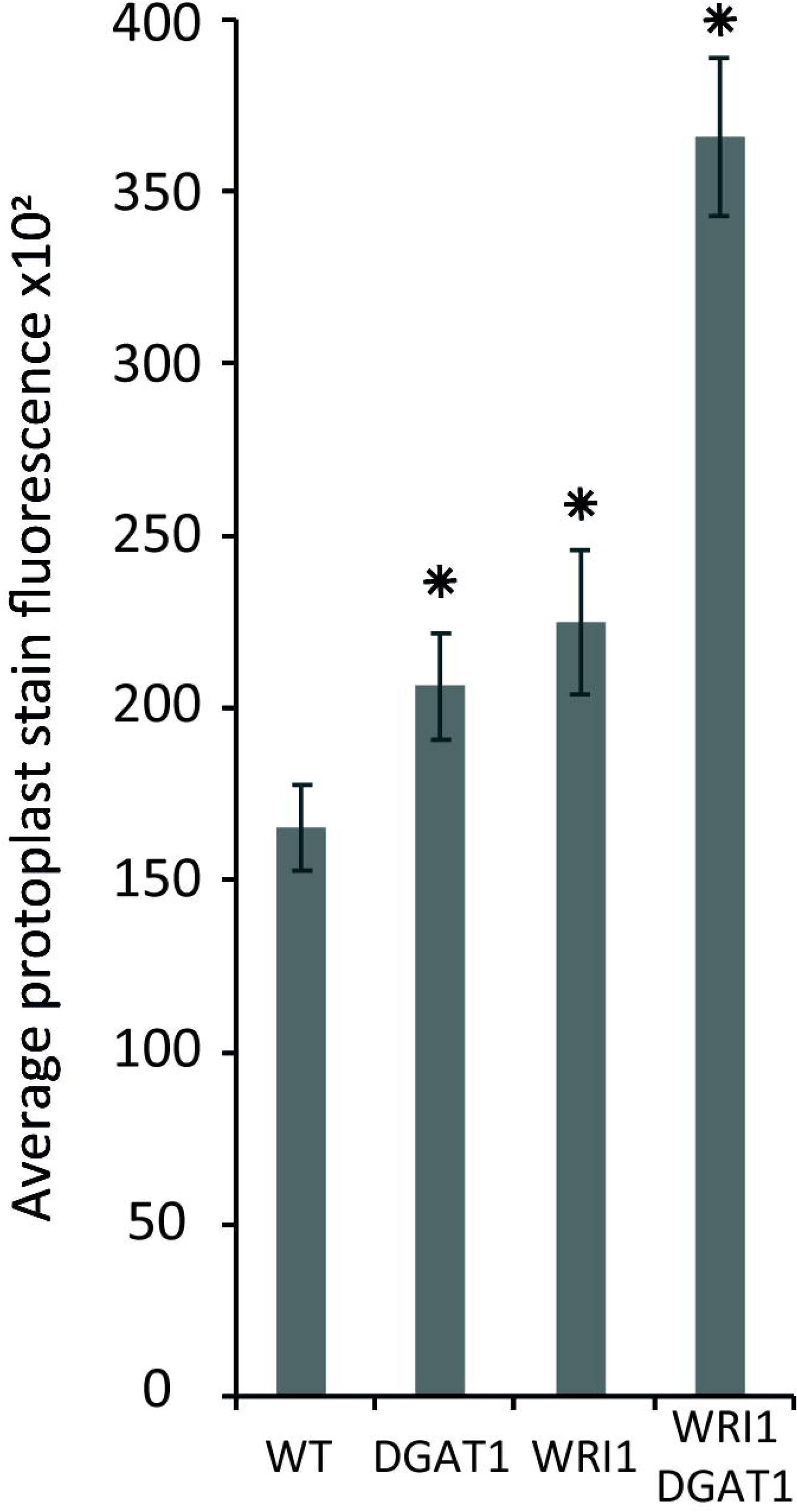
Lipid accumulation in tobacco leaf transiently transformed protoplasts. Lipid accumulation WT tobacco leaf protoplasts compared to WT protoplasts transiently transformed with *WRI1* or *DGAT1*, was measured with BODIPY staining and flow cytometry recording of the average particle stain fluorescence for the whole sample. Error bars represent the standard deviation. Asterisks represent statistically significant differences (P value<0.05) compared with WT.

**Figure 5.**
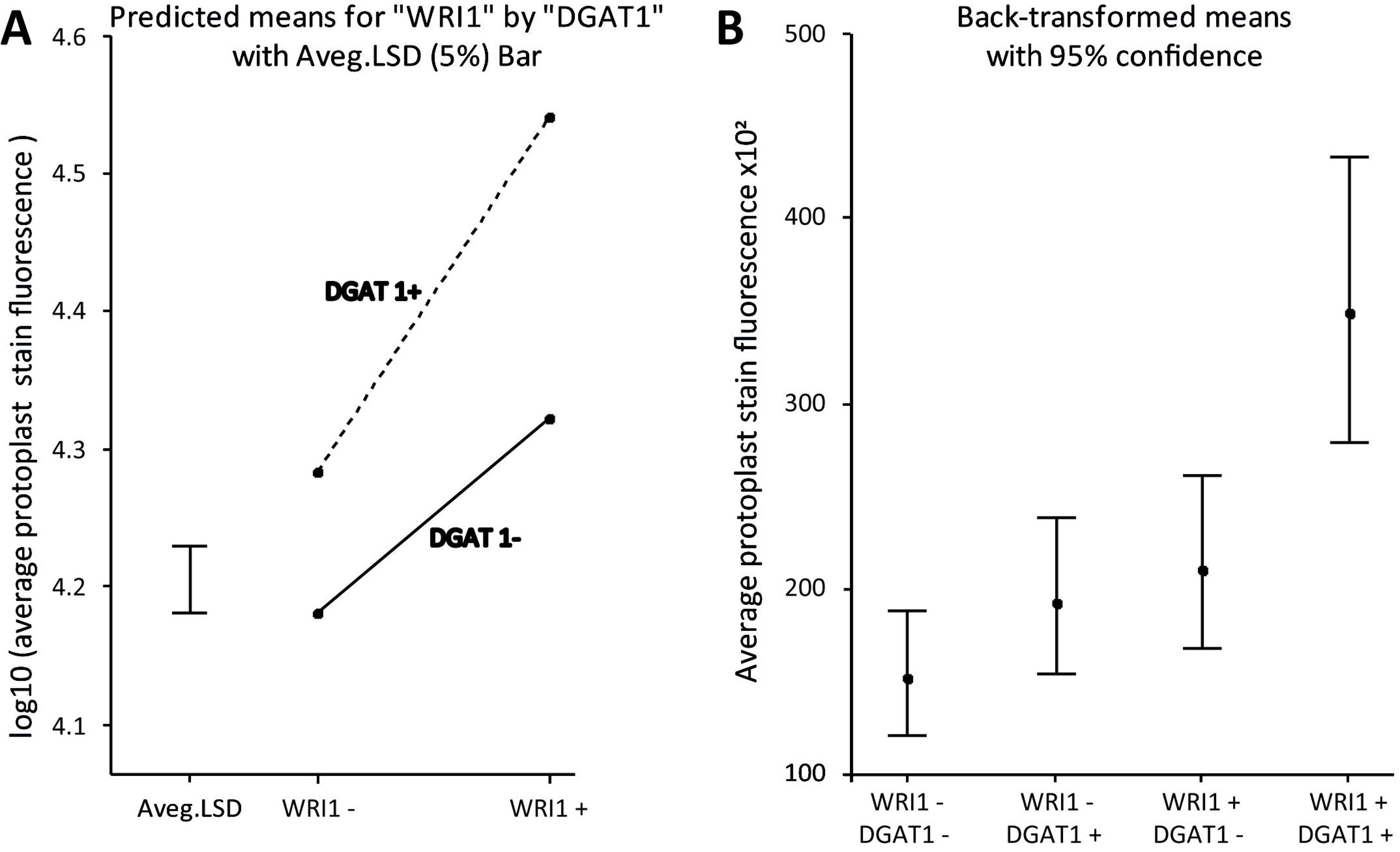
*WRI1* and *DGAT1* have a synergistic positive effect on protoplast lipid biosynthesis. **(A)**, Interaction plot of predicted treatment means with 5% Least Significant Difference (LSD) bar on the log10-transformed scale and **(B)**, plot of the back-transformed predicted means (with back-transformed 95% confidence interval limits) from the analysis of the data previously presented in Figure 4B. WT corresponds to “WRI1-:DGAT1-” and the samples transformed with *DGAT1, WRI1* and both, correspond to WRI1-:DGAT1+, “WRI1+:DGAT1-” and “WRI1+:DGAT1+” respectively.

In fitting any statistical model, it is important to assess various residual diagnostic plots to ensure that the assumptions underlying the model are met and that predictions and tests derived from the model are reliable. Residual diagnostic plots from the fit of the model to the raw data suggested that the random scatter in the data increases as the mean of the data increases; a common property of such assay data. To model the data appropriately, a log10 (log to base 10) transformation of the data was required to ensure that the model assumption of constant variance was met. The statistical model was therefore applied to the log10 transformed data, and predicted means were back-transformed as required to obtain predictions on the original data scale. Treatment predicted means and associated statistics were obtained using the R ‘predictmeans’ package (Luo et al., 2018). Tests for the statistical significance of terms in the factorial treatment structure were obtained using the R ‘lmerTest’ (Kuznetsova et al., 2017) package.

The errors bars and P values presented in figures 2, 3, 4 and 6 were calculated using respectively the Excel functions STDEV and T.TEST (array1, array2, tails=2, type=3).

**Figure 6.**
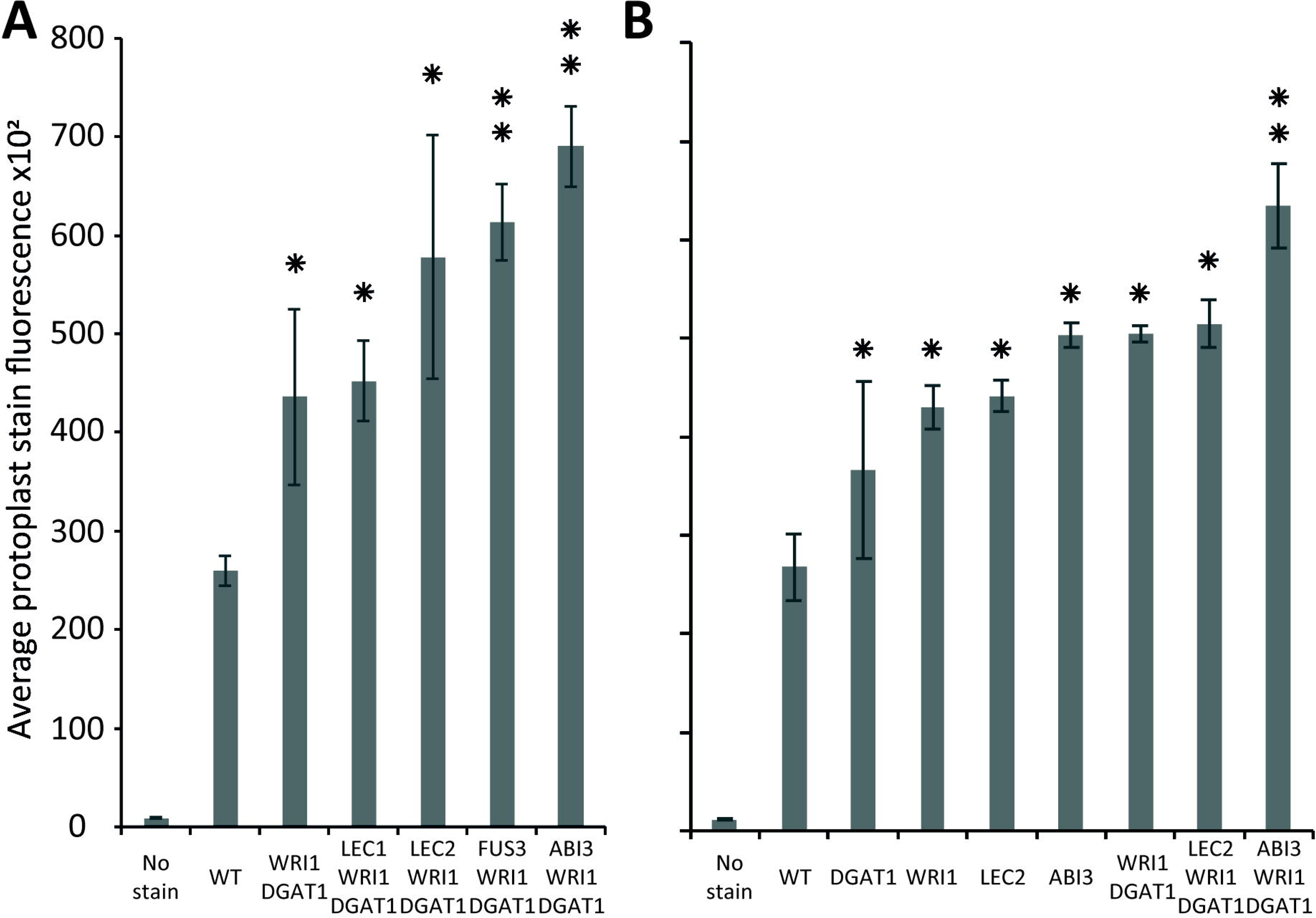
Comparison of the relative effect of various transiently transformed effectors on lipid accumulation in tobacco leaf protoplasts. Lipid accumulation, compared to WT, in tobacco leaf protoplasts transiently transformed with *WRI1, DGAT1, ABI3, FUS3, LEC1* or *LEC2* and various combinations, was measured by flow cytometry and based on the average particle stain fluorescence for the whole sample following BODIPY staining. Bars represent standard deviation. **(A)**, Comparative additional effect of *ABI3, FUS3, LEC1* or *LEC2* on lipid content in protoplasts co-transformed with *WRI1* and *DGAT1*. **(B)**, Individual effect of *WRI1, DGAT1, ABI3, LEC2* and of various combinations on lipid content in co-transformed protoplasts. A single star represents statistically significant difference (P value<0.05) compared with WT, while the double stars represent additional statistically significant difference (P value<0.05) compared with *WRI1* and *DGAT1* combination, (shown for samples with significant increase only).

## Results

### A new and rapid method for testing DNA constructs for plant lipid engineering

The new protoplast based-platform for high throughput plant lipid engineering presented in this study allows for the testing of DNA constructs about their ability to induce lipid accumulation in plant cells within only 72 hours. Using BODIPY™ 493/503, a fluorescent neutral-lipid stain, lipid quantity in protoplasts is evaluated by measuring cell fluorescence by flow cytometry or alternatively by fluorescence-activated cell sorting if specific cell populations need to be separated for further analyses (**figure 1**). Protoplasts with high lipid accumulation consequently exhibit higher fluorescence and can be identified and eventually sorted based on the relative intensity of the fluorescent stain per cell.

### Protoplasts can be used as a predictive tool for plant lipid engineering

To confirm that protoplasts can be used as a predictive tool for plant lipid engineering, we tested if the high oil phenotype of a previously established stably transformed line could be reproduced with protoplasts isolated from the same line. The surprising lack of studies regarding lipid accumulation in protoplasts to date suggested that such approaches might not be possible to be conducted. We previously reported the establishment of a High-Oil line of tobacco (henceforth referred to as HO) accumulating in its leaves up to 24 times more total lipid, and particularly 76 times more triglycerides, at maturity, compared to wild-type (henceforth referred to as WT) (Vanhercke et al., 2014). We also reported that this increased lipid accumulation was gradual during development, thus potentially making it difficult to measure the increase in very young tissue. Leaf tissues from 15 days old WT and HO plants were harvested and partly used for protoplast isolation. The remaining leaf tissues as well as protoplasts isolated from the same leaf samples were then analysed for their triglycerides and total lipid content (**figure 2**). The analyses revealed that at this early developmental stage and under the specific growth conditions of this assay, HO plantlets had accumulated about two times more total lipid and almost 90 times more triglycerides than WT, while HO protoplasts had accumulated about 25% more total lipid and more than two times more triglycerides than WT. Thus, despite some variation in intensity, the analyses confirmed that the increase can still be measured at the protoplast level. We also confirmed that even though the variation between the HO line and WT would be easier to measure by focusing on the triglycerides level, it is still quite feasible to measure it at the total lipid level. This will allow for the use of generalist neutral lipid stains, which stain any neutral lipid and not only triglycerides, to measure the variation in oil content.

### Tobacco leaf protoplasts can be sorted based on lipid content

In the previous part, we demonstrated that we can measure variation in protoplast oil accumulation by conducting the analysis at the total lipid level instead of focusing on triglyceride accumulation only, which should allow for the use of a generalist neutral lipid stain, which stains triglycerides and all other neutral lipids, to measure the variation in oil content. We next sought to confirm that protoplasts can be sorted based on storage lipid content. Protoplasts were isolated from the leaves of 15-day-old WT and HO plantlets as described above, and stained with BODIPY™ 493/503, a generalist neutral lipid stain, prior to microscopy (**figure 3A**) or flow cytometric analysis (**figure 3B**). WT and HO stained protoplasts were consecutively sorted, based on their intensity of BODIPY™ 493/503 fluorescence relative to chlorophyll auto-fluorescence. Previous iterations of sorting coupled with microscopy allowed us to determine where the intact green cells are situated in the graph generated during cell sorting (data not shown). Two distinct populations of intact green cells were gated, with P1 defined as the most intensively stained cells and P2 as the rest of the intact green cells (figure 3B). Average stain fluorescence per cell was calculated for each population (**figure 3C left**). For each population, three samples of 250,000 cells were collected and then subjected to the same total lipid analysis as described above (**figure 3C right**). This was not done for the WT P1 population because it was not possible to retrieve enough cells to conduct the analysis. Several conclusions could be drawn for this assay.

First, we confirmed that BODIPY™ 493/503 staining allows for measuring of the oil content difference that we previously measured between WT and HO protoplasts. The flow cytometric analysis showed that the HO sample had a distinct P1 population that segregated from the rest of the cells, while the WT cells attributed to the P1 population corresponded to the top cells of the P2 population rather than being a distinct cell population (**figure 3B**). For both WT and HO samples, cells of the P1 population exhibited higher average stain fluorescence intensity compared to cells of the P2 population (**figure 3C left**). Lipid analyses revealed that the HO-P2 population was significantly more lipid-rich than the WT-P2 population and that the HO-P1 population was significantly more lipid-rich than both the HO-P2 and WT-P2 populations (**figure 3C right**). These results correlate with the corresponding measurements of stain fluorescence intensity for each population (**figure 3C left**), illustrating the direct correlation between lipid content and BODIPY™ 493/503 fluorescence intensity. Secondly, the HO whole sample had a higher average protoplast stain fluorescence intensity than the WT whole sample (green bars in **figure 3C left**). This correlates with the results presented in the previous part and demonstrates that HO protoplasts accumulate more total lipid than WT.

Overall, this assay confirmed that protoplasts can be sorted based on lipid content and that measuring the whole sample BODIPY™ 493/503 fluorescence signal (green bars in **figure 3C left**) is enough to evaluate the effect of a gene combination on the overall lipid content in protoplasts. We also note that the variation measured between the WT and HO sample is amplified when using BODIPY™ 493/503 staining coupled with flow cytometry (compared with lipid analysis methods), suggesting that this alternative screening strategy could be more powerful than traditional ones.

### Transiently transformed tobacco leaf protoplasts can accumulate lipids

We have established that leaf protoplasts from a stable transgenic line that accumulates more oil in its leaves than WT also accumulates on average more oil per protoplast compared to wild type and that the difference can be measured by traditional lipid analyses or measured by flow cytometry after protoplast isolation and appropriate staining. However, to date, it was unclear if and how leaf protoplasts would produce an increased quantity of oil compared to WT, after isolation and transient transformation, reflected by the absence of any similar study in the literature.

The stable transgenic line used in the study over-expresses a combination of genes coding for the *Arabidopsis thaliana* WRI1 transcription factor, a regulator of the late glycolysis and de novo fatty acid biosynthesis pathways, the *Arabidopsis thaliana* DGAT1 acyl-CoA: diacylglycerol acyltransferase, a major lipid metabolism enzyme, and an OLEOSIN oil body protein from *Sesamum indicum*, which coats oil droplets and protects them from degradation. The reasoning for including OLEOSIN was that achieving greater oil accumulation in leaves in a long gene-expression-window (stable expression) might have required reducing oil degradation in this non-seed tissue. However, we previously demonstrated that the combined transient overexpression of *WRI1* and *DGAT1* only in *Nicotiana benthamiana* leaves was enough to produce a measurable and significant synergistic effect on leaf oil content (Vanhercke et al., 2013).

Based on the results presented above, next we tested transient transformation of leaf protoplasts with genetic constructs carrying expression cassettes for *WRI1, DGAT1* or both and measured the difference in protoplast total neutral lipid content, using BODIPY™ 493/503 staining and flow cytometry recording of the average protoplast stain fluorescence for the whole sample (**figure 4**). Compared to the non-transformed control, leaf protoplasts transiently transformed with *WRI1, DGAT1* or both exhibited increased average stain fluorescence, indicating an increase in total neutral lipid content.

We also observed that the synergistic effect of *WRI1* and *DGAT1* co-transformation, previously reported in leaf transient assays, was noticeable and significant in our protoplast assay, which is further discussed below.

### Combinatorial gene testing in protoplasts mimics results obtained with leaf transient assay

The synergistic effect of *WRI1* and *DGAT1* on tobacco leaf protoplast oil content when co-overexpressed, compared with their individual effect when separately over-expressed, and previously reported with the *Nicotiana benthamiana* leaf transient assay (Vanhercke et al., 2013) and in stably transformed *Nicotiana tabacum* plants (Vanhercke et al., 2014), was also observed in our protoplast assay. Indeed, as we demonstrated, the effect of each gene or gene combination on the lipid accumulation trait of a protoplast population transformed with it, can be assessed by measuring the increase of fluorescence of this population, compared to a WT population, after BODIPY™ 493/503 staining. Our results suggest that when co-transformed, the combined effect of *WRI1* and *DGAT1* on lipid accumulation is greater than the additive effect of each individual transformations, illustrating the synergistic effect of *WRI1* and *DGAT1* (**figure 4**).

The statistical analysis of the data confirms the statistical significance of the synergistic effect, with a p-value for the interaction effect on the log10-transformed scale of p < 5e-6. An interaction plot of the WRI1 by DGAT1 predicted means is given in **Figure 5A**. The significant p-value confirms that the lack of parallelism of the lines (and hence the presence of the synergistic effect) joining the pairs of means is genuine. **Figure 5B** plots the predicted means back-transformed to the scale of the data, with associated back-transformed 95% confidence interval limits for the means indicated by the error bars.

Overall our results confirm that protoplast systems are robust models that can be used to predict and inform plant phenotype and are compatible with combinatorial gene testing. This finding is particularly important for coupling transient protoplast expression with high throughput combinatorial gene screening.

### ABI3 induces lipid accumulation in tobacco leaf protoplasts

Our new screening strategy constitutes a new high throughput tool to fast-track plant lipid engineering strategies and offers an alternative method to study plant lipid metabolism. We therefore wanted to use this newly developed strategy to investigate the relative effect of the major known seed specific developmental regulators. ABI3, FUS3, LEC1 and LEC2 are master regulators that control the gene regulation networks governing most seed developmental mechanisms. FUS3, LEC1 and LEC2 also have been reported to trigger oil accumulation through regulation of *WRI1*, which controls the expression of many genes essential for fatty acid synthesis. However, while many targets of ABI3 were identified to function in seed oil storage, the direct effect of *ABI3* on lipid content in plants remained undocumented.

Tobacco leaf protoplasts were transiently transformed with *WRI1* and *DGAT1* only or with one of the master regulators, being *ABI3, FUS3, LEC1* or *LEC2*. The effect on protoplast total neutral lipid content was then measured and compared as described above (**figure 6A**). This first assay indicated that in addition to the combined effect of *WRI1* and *DGAT1*, only co-expression with *FUS3* and *ABI3* significantly (P value < 0.05) further increased protoplast total neutral lipid content. This was particularly surprising due to the lack of a previously reported direct effect of *ABI3* overexpression on plant lipid accumulation. To date, the most efficient gene combination reported for lipid accumulation enhancement involved *WRI1* with *DGAT1* and *LEC2* or *SDP1* silencing (Vanhercke et al., 2017). Therefore, in a second similar analysis we investigated in more detail the individual effect of each individual gene (*WRI1, DGAT1, LEC2* and *ABI3*) and of several related combinations on total neutral lipid content in transiently transformed protoplasts (**figure 6B**). This second assay confirmed that *ABI3* overexpression alone resulted in the highest accumulation of total neutral lipids in protoplasts. In addition, unlike *LEC2*, co-expression with *ABI3* further increased the lipid content of protoplasts transiently transformed with *WRI1* and *DGAT1*. This difference supports the previous reports that *LEC2* regulates fatty acid biosynthesis mainly through the regulation of *WRI1* (Baud et al., 2007; Santos-Mendoza et al., 2008). Thus, the addition of *LEC2* might be ineffective due to the expression level of exogenous *WRI1* in the co-transformed protoplasts being already so high that triggering the expression of native *WRI1* would make a negligible effect on the overall *WRI1* expression levels in protoplasts. Overall, our results suggest that *ABI3* control over plant lipid metabolism and storage might involve mechanisms that act independently from the previously described *LEC2* and *WRI1* regulation networks. Our new screening approach also suggests that *ABI3* could be an important factor triggering oil accumulation in plants and suppose that *ABI3* should be of prime interest in future plant lipid engineering approaches.

## Discussion

### Transient protoplast transformation as an alternative predictive tool for plant lipid engineering

In this study, we demonstrated that protoplasts from 15-day-old tobacco leaves are a reliable model to study plant lipid engineering. This protoplast-based screening assay takes about three days to complete, as compared to the minimum of two weeks required for a traditional transient leaf transformation assays based on agroinfiltration and GC analysis. We confirmed the direct correlation between protoplast lipid content and protoplast fluorescence intensity after lipid specific fluorescent staining. Furthermore, we show that protoplasts can be sorted based on lipid content and further used for downstream processing and lipid or molecular analyses. Our results demonstrate that transient gene expression in protoplasts produces lipid accumulation results similar to those obtained in stably transformed plants. We confirm that combinatorial gene testing is possible in protoplasts, with synergistic effect of the genes *WRI1* and *DGAT1* on increasing lipid accumulation in protoplasts, comparable to previously published studies conducted in transiently and stably transformed leaves (Vanhercke et al., 2014; Vanhercke et al., 2013).

### Highlight on ABI3 as a new major lead for improvement on lipid accumulation in crops

Using our newly developed workflow, we demonstrated that *ABI3* has a significant effect on protoplast total neutral lipid content. *ABI3* super-transformation significantly further increased the lipid content of protoplasts transiently transformed with *WRI1* and *DGAT1*, to a higher level than other known major effectors.

This result suggests that *ABI3* control over plant lipid metabolism and storage might at least partially involve mechanisms that are independent from the previously described *LEC2* and *WRI1* regulation networks. Our new approach also suggests that *ABI3* is a major regulator of oil accumulation in plants and reveal ABI3 as a new target for future plant lipid engineering approaches.

### A new high throughput platform for gene function and plant metabolic engineering study

The protoplast-based workflow presented in this study is very versatile and should be compatible with many crop species and for a wide range of future output traits. Indeed, protoplasts can nowadays be isolated from almost any plants and any tissue. A profusion of methods were developed during the 70’s and 80’s for many major crops and model plants, including tobacco (*Nicotiana tabacum*) (Guerche et al., 1987a; Takebe and Otsuki, 1969), cowpea (*Vigna unguiculata*) (Hibi et al., 1975), tomato (*Solanum lycopersicum*) (Motoyoshi and Oshima, 1979), turnip (*Brassica rapa*) (Howell and Hull, 1978), barley (*Hordeum vulgare*) (Loesch-Fries and Hall, 1980), cucumber (*Cucumis sativus*) (Maule et al., 1980) great millet (*Sorghum bicolor*), rice (*Oryza sativa*), wheat (*Triticum monococcum*) (Bart et al., 2006; Ou-Lee et al., 1986), canola (*Brassica napus*) (Guerche et al., 1987b), *Arabidopsis thaliana* (Damm et al., 1989), and maize (*Zea mays*) (Sheen, 1990). Such approaches then fell into disuse probably due to the very high technical skills required to perform high quality protoplast isolation, transformation and culture. However, new methods were sporadically released for other crops of interest, such as carrot (*Daucus carota*) (Liu et al., 1994). Recently, in the wake of a multitude of new high throughput cutting-edge technologies becoming easily accessible, protoplast-based systems have regained in their popularity and a profusion of improved methods have been released. Protoplast systems are the most suited for rapid screening of gene silencing (siRNA and miRNA) and genome-editing (e.g. CRISPR) targets (Cao et al., 2014; Maćkowska et al., 2014; Schapire and Lois, 2016; You et al., 2014; Zhai et al., 2009). In addition, protoplast-based systems are now available for almost any plant of commercial interest, including lettuce (*Lactuca sativa*) (Sasamoto and Ashihara, 2014), grape (*Vitis vinifera*) (Wang et al., 2015) and bean (*Phaseolus vulgaris*) (Nanjareddy et al., 2016) and even the less expected, such as oil palm (*Elaeis guineensis*) (Masani et al., 2014), poplar (*Populus euphratica*) (Guo et al., 2015) and switchgrass (*Panicum virgatum*) (Burris et al., 2016).

In addition, our strategy should allow similar approaches for a broad variety of other metabolic traits since many fluorescent dyes, specific to particular metabolites are available. In the absence of a specific stain, genetic switches or biosensors can be developed for the specific detection of small molecules such as toluene (Behzadian et al., 2011), metabolites (Liu et al., 2017), and even specific proteins of interest (Abe et al., 2011; Wongso et al., 2017). For example, a new phosphatidic acid (PA) biosensor was recently developed to track PA cellular concentration and dynamics (Li et al., 2019). PA is the precursor of most phospholipids and triglycerides and represents around 1% of all lipids. PA acts as a key signalling lipid that predominantly accumulates at the plasma membrane. This biosensor was based on the use of Förster resonance energy transfer (FRET) and was used to study the spatio-temporal complexity of PA accumulation in plant tissues and its contribution to mediate plant response to salt stress. Numerous other biosensors have been also developed based on FRET to study protein-protein, protein-DNA or DNA-DNA interactions (Crawford et al., 2012; Lymperopoulos et al., 2010). Alternatively, riboswitches have been engineered as nucleotide-based biosensors with great potential (Hallberg et al., 2017; Villa et al., 2018).

The new screening strategy presented in this study constitute the first step towards high throughput screening of complex genetic libraries in a plant system by using transient protoplast transformation and rapid cell sorting based on a fluorescent signal that is linked to a trait of interest. The workflow can be completed as a single experiment in a matter of days as opposed to several years that are needed by conventional means. Such novel synthetic biology-inspired high throughput screens are paving the way for rapid identification of new unreported genes or gene combinations that significantly improve traits of interest in crops and thereby greatly improve the timelines associated with crop improvement (Zhang et al., 2019).

## Authors contribution

Benjamin Pouvreau, Thomas Vanhercke and Surinder Singh conceived the study; Benjamin Pouvreau and Cheryl Blundell performed experimental work with assistance from Harpreet Vohra for fluorescence-activated cell sorting; Benjamin Pouvreau analysed the data with assistance from Alec Zwart for statistical analysis; Benjamin Pouvreau wrote the manuscript with contributions from all authors.

## Acknowledgments

Thanks to Nathalie Niesner for cloning the *DGAT1* fragment into the “CaMV35S promoter / NOS terminator” expression cassette of the modified pENTR11 vector.

Thanks to Lauren Venugoban for aiding with microscopy.

This work was partly funded through the CSIRO Synthetic Biology Future Science Platform and the CSIRO Research Office CERC Postdoctoral Fellowship scheme.

